# On branch lengths assignment methods for trees with fixed topology and related biological applications

**DOI:** 10.1101/2024.07.29.605688

**Authors:** Wei Wei, David Koslicki

## Abstract

Distance-guided tree construction with unknown tree topology and branch lengths has been a long studied problem. In contrast, distance-guided branch lengths assignment with fixed tree topology has not yet been systematically investigated, despite having significant applications. In this paper, we provide a formal mathematical formulation of this problem and propose two representative methods for solving this problem, each with its own strength. We evaluate the performance of these two methods under various settings using simulated data, providing guidance for the choice of methods in respective cases. We demonstrate a practical application of this operation through an extension we termed FunUniFrac, which quantifies the differences in functional units between metagenomic samples over a functional tree with assigned branch lengths, allowing clustering of metagenomic samples by functional similarity instead of taxonomic similarity in traditional methods, thus expanding the realm of comparative studies in metagenomics.

## I. Introduction

**T**REE construction has been a commonly encountered problem in biology. One such example is the construction of phylogenetic trees in the study of evolution. A phylogenetic tree is a weighted binary tree that reflects the evolutionary relationship among species, in which the organisms are represented as nodes in the tree and the evolutionary proximity between any pair of organisms represented by the length of the path that connects them. Often times, the exact evolutionary trajectory is unknown, both in terms of branching points and edge lengths. Such information will then need to be constructed based on biological data such as sequence alignments. This kind of tree construction involving the establishment of tree topology and edge lengths concurrently using leaf-level information has been a widely studied problem since as early as the 1960s [1], oftentimes in the context of phylogenetic tree construction. Since then, many methods have been proposed attempting to solve this problem. These methods were generally thought to fall into one of the two broad categories: distance-based and character-based. The former makes use of a distance matrix while the latter considers a number of probable trees and chooses one that is most likely according to evolutionary assumptions [2]–[Some of the most commonly used distance-based methods include Neighbor Joining (NJ) [5] and unweighted pair group method using arithmetic averages (UPGMA) hierarchical clustering. These methods start off from a distance matrix and find ways to successively connect the nodes to form internal nodes, constructing tree topology and branch lengths at the same time. Representatives of character-based methods include the maximum likelihood method [6] and the maximum parsimony method [7]. The former exhaustively considers all possible trees and uses probabilistic models to pick the tree that is the most probable, whereas the latter considers only the informative sites of the multiple sequence alignments and selects the tree that minimizes the number of mutations.

The construction of these tree structures is without a doubt meaningful and has contributed significantly to computational biology. Not only does it allow us to have more insight on the hierarchical relationship among organisms, but it also provides the structure for tree-based analyses and computations to be applied on biological data that previously only existed in the form of individual sequences. An example of such analyses is the computation of the UniFrac metric [8] in the differential abundance analysis of metagenomic samples. In the computation of UniFrac, metagenomic samples are represented as vectors indexed by microorganisms present in all samples, with each entry of a vector representing the relative abundance of the organism in that sample. In addition, a phylogenetic tree, either pre-existing or constructed ad hoc, consisting of all microorganisms in the samples, is required. UniFrac then computes Earth Mover Distance over a phylogenetic tree to quantify sample dissimilarity [9]. The UniFrac metric, known as a phylogenetic beta-diversity metric due to its incorporation of phylogenetic information, is shown to be more robust in some cases than its non-phylogenetic counterparts such as the Jaccard index or Bray-Curtis dissimilarity [10]. This superiority of phylogenetic-based analysis demonstrates the importance of structural information, both in terms of topological structure and numerical values of edge lengths; the absence of either component would deem the computation of UniFrac impossible.

In the scenarios of tree construction described above, the topological information and branch lengths are intertwined, both dependent on leaf-level sequence information. There exist, however, many other types of tree-like structures both in and outside of the biological realm with a topology not entirely dependent on readily comparable traits such as sequences.

Examples of such biological structures can include taxonomic trees in which species are classified based on similarities in genomic and phenotypic traits, biological behaviors and ecological habitats; as well as metabolic pathways where components are grouped together due to involvement in related biochemical processes. In a broader context, there are also decision trees in the context of machine learning, social networks, knowledge graphs or concept hierarchies.

In some cases, weights can be assigned to edges *a posteriori* using experimental data, such as in the case of metabolic pathways. In other cases, those structures simply do not come with natural branch lengths, such as in the case of taxonomic trees or knowledge graphs. In these cases, edges often simply represent inclusion or some kind of non-numerical relation. However, even for structures without naturally defined branch lengths, having some kind of weight on branches can be largely beneficial, as it allows graph-based algorithms to be applied on those structures in analyses. An example is WGSUniFrac [11], which involves replacing a phylogenetic tree with a taxonomic tree followed by assigning branch lengths, thus allowing UniFrac to be computed on whole genome shotgun data, which was previously impractical. Like this, assigning branch lengths to a pre-existing tree opens the door for analyses to be performed on a wider range of data and expands the utility of existing methods.

In contrast to distance-based phylogenetic tree construction, imputing branch lengths of a generic tree structure with a fixed topology has been a less frequently studied area. In many cases, it is only considered as a sub-task of tree constructions in which the real topology is to be determined. For example, in both the works of Fitch & Margoliash [1] and Pardi & Gascuel [12], branch lengths of an “initial tree” with an assumed topology are first computed based on the distance matrix, followed by either the search of alternative trees or adjustment of the initial topology to determine the optimal topology that best agrees with the given distance matrix.

However, imputing branch lengths of a fixed topology is by no means a less important problem. On the contrary, it is a problem with a wider scope of potential application. Though previous works such as Fitch & Margoliash [1], Pardi & Gascuel [12] (who considered only binary trees) and Farris [13] have laid the foundation for such a case such as least-square minimization [4] and Farris transform [13], is is only with the context of phylogeny inference (eg. with assumptions about character-based or evolutionary model-based distances; inferring/constructing the topology, etc.). To the best of our knowledge, there has not been a formal and rigorous examination of this generic mathematical problem with generic but fixed topology and generic distance metric, nor is there a thorough analysis of the behavior of the methods under different conditions.

In this study, we explored the mathematical backbone of the problem of imputing branch lengths of a fixed tree topology given leaf-level pairwise distances. We proved the mathematical conditions for the solvability of this problem and proposed two classes of methods to solve it, including an efficient implementation of an adopted version of the traditional Farris transform. Through theoretical and experimental explorations, we evaluated the performance of these two classes of methods under different scenarios and demonstrated a real-life application of this operation by imputing the branch lengths of the KEGG [14]–[16] hierarchy using estimated distances among orthologous proteins. We further developed a new metric which we termed the FunUniFrac, inspired by the UniFrac metric [8], which, coupled with the KEGG hierarchy with assigned branch lengths, was able to compute the functional differences between metagenomic samples. The code for the experiments done in this paper can be found here: https://github.com/KoslickiLab/branch-lengths-assignment.git.

## II. Problem and methods

### A. Problem and scope

We begin by giving a formal mathematical description to this problem. Suppose we are given a tree *T* with edge set *E* of size *n* such that every *e*_*i*_ in *E* has a non-negative length. Let *m* be the number of leaf nodes in *T* and *D*_*T*_ be an *m ×m* matrix representing the pairwise tree/geodesic distances among the leaf nodes of *T*. We call *D*_*T*_ the distance matrix associated with *T*. The relationship between *T* and *D*_*T*_ can then be represented in the following way. Let *x* be a vector of length *n* such that *x*_*i*_ is the length of *e*_*i*_. Let *y* be a vector of length 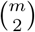 consisting of values in *D*_*T*_ in an arbitrary order, any let *A* be a binary matrix of size 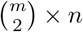 such that given any index 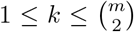 of vector *y*, if *y* = *D* (*i, j*) where *i* and *j* leaf nodes in *T*, then the *k*th row of *A, A*(*k*, ·), is a binary representation of the edges involved in the path between leaf nodes *i* to *j*. i.e.

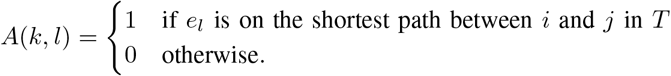

We call *y* the distance vector derived from *D*_*T*_. Then we have the linear system

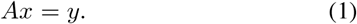

Conversely, if we are given the topology of a tree *T* and *D*_*T*_, solving for *x* in this linear system gives the lengths of the edges in *T*. This procedure can be computationally intensive in real-world biological applications. For instance, if *T* is a full binary tree as in the case of most phylogenetic trees, if *T* has a height *h*, then 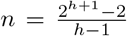 and *m* = 2^*h*^, and as a result, *A* would have a dimension of 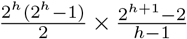, with significantly many more rows than columns. In other words, the linear system (1) would be overdetermined and the solution for *x* will now depend on the properties of *T* and *D*_*T*_ and the relationship between them.

Our objective is to recover *x* given *T* and *D*_*T*_. To examine the quality of this solution under different conditions, we first place some restrictions on *T* by defining the following properties.

#### Definition II.1

(Compatible distance matrix with respect to a given tree). Given a tree *T* with edge set *E*, we let *T*_*ij*_ denote the collection of edges in the shortest path between nodes *i, j* ∈ *T*. For a given distance matrix *D* indexed by all or some of the the nodes in *T*, if there exists a mapping *f* : *E* → ℝ such that for every pair *x* and *y* in the indices of *D*, we have 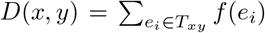, we say that *D* is compatible with *T*. Otherwise, we say *D* is incompatible with *T*. We call *f* an assignment function for the edges of *T*.

**Remark**. Another way to look at compatibility is that if a matrix *D*_*T*_ is not compatible with *T*, then attempting to solve the edge lengths of *T* will result in an inconsistent solution. An example of a compatible matrix and an incompatible matrix is shown in Supplementary Materials Section 1. Obviously, if all the edges in *T*_*ij*_ have known branch lengths and we construct *D*_*T*_ such that *D*_*T*_ (*i, j*) is precisely the sum of lengths of all edges in *T*_*ij*_ for all *i, j* in the indices of *D*_*T*_, then *D*_*T*_ will be compatible with *T*. In this case *f* maps each edge in *E* to its original known length.

There are two main factors that impact the compatibility of a distance matrix with a tree. The first being that the distance measurement is prone to errors. The second being that the nature of the distance measurement metric not being reflected in the tree topology. For example, two nodes may be siblings in the tree but have a very large distance in the distance matrix, perhaps even larger than the distance between some of their descendants. Based on the results shown in Figure 5, we conjecture that the second case is more detrimental than the first in increasing the difficulty in reconstructing the branch lengths.

It is important to distinguish between compatibility defined above from the concept of “additivity” frequently discussed with respect to a given distance matrix, but often in the context of unfixed tree topology. In fact, compatibility is a stronger condition than additivity. A detailed explanation of the distinction between additivity and compatibility is discussed in Supplementary Section 1.2 with a counter-example demonstrating how a distance matrix can be additive and yet incompatible with a given tree. Essentially, additivity ensures that there exists a tree that is compatible with a given matrix, whereas compatibility describes the relationship between a fixed tree and a fixed distance matrix.

If *D*_*T*_ is compatible with *T*, an assignment function *f* can be found. However, this assignment function may not be unique. For *f* to be unique, we need the following property of the tree to hold true.

#### Definition II.2

(Unambiguous tree). We say that a tree *T* is unambiguous if the following conditions hold true:

1. Every node in *T* except the root node has at least one sibling.
2. The root node has at least three children.

We will show in the next section that if *T* is unambiguous and *D*_*T*_ compatible with *T*, a unique assignment function *f* can be found. For the sake of clarity in understanding this proof, we clarify the concept of level, which sometimes has variable meanings depending on context, within the scope of this paper as follows.

#### Definition II.3

(Level). The level of a node in a tree is defined to be the number of edges between the node and the root. The level of a tree is the level of the node that has the maximum level in the tree.

### B. Solving the problem in the perfect scenario

In the most ideal scenario, *x* is recovered exactly as it is. In this section, we show the conditions for this to take place and provide an algorithm to achieve this objective.

#### Lemma 1.

Given any one-level tree with more than two leaf nodes, the associated matrix *A* in linear system 1 is a full-rank matrix.

*Proof*. Suppose the given tree has *m* leaf nodes where *m* ≥ 3. We show that the associated matrix *A* has rank *m* by construction. Suppose the *m* leaves are labeled *n*_1_ through *n*_*m*_ with the lengths of edges leading to the nodes labeled *x*_1_ through *x*_*m*_ respectively. Consider the following linear system:

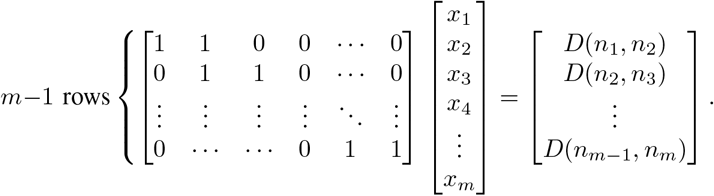

We can make the following observations. First, the matrix on the left with *m* − 1 rows is a sub-matrix of *A*. Also, it can be easily checked that the *m* − 1 rows are linearly independent, giving *A* a rank of at least *m* − 1. We claim that we can find another row in *A* that is linearly independent with the *m* − 1 rows, giving *A* a rank of *m*. Case 1: *m* is even. In this case, we claim that the vector (1, 0, …, 1, 0) is linearly independent to all the *m* − 1 row vectors in the sub-matrix of *A* above. Note that this vector is indeed from matrix *A* because it corresponds to the distance *D*(*n*_1_, *n*_*m* −1_). Appending this row to the last row of the *m* − 1-row sub-matrix of *A*, we obtained a a *m*-row sub-matrix of *A*, which we will call *Â*. We now show that *Â* has rank *k* by performing Gaussian Elimination. Subtracting row *m* by row 1 turns row *m* into (0, − 1, 0, …, 1, 0). Next, add row 2 to row *m*, turning row *m* into (0, 0, 1, 0, …, 1, 0). Repeating this process of successively subtracting odd rows from row *m* and adding even rows to row *m* eventually leaves row *m* in the form of (0, 0, …, − 2), which is linearly independent to all the other *m* − 1 rows.

Case 2: *m* is odd. We claim that (1, 0, …, 0, 1) corresponding to *D*(*n*_1_, *n*_*m*_) is linearly independent to all the other *m* − 1 vectors. Performing the same operation, we will obtain (0, 0, …, 0, 2) as the last row of *Â*, giving *Â* rank *m*. Since *Â* is a sub-matrix of *A*, we conclude that *A* also has rank *m*.

This proof is valid for any *m* ≥ 3. This also shows that for an unambiguous tree with one level, the linear system (1) has a unique solution.□

#### Claim 1.

In the event where *T* is unambiguous and *D*_*T*_ is compatible with *T*, the linear system (1) has a unique solution.

*Proof*. By Lemma 1, this holds true in the case where *T* has only one level. We now move on to the general case where *T* has more than one level. We begin by showing that all the leaf edges (edges leading to a leaf node) have a unique assignment of length.

Suppose for the sake of contradiction that there are at least two assignment functions satisfying *D*_*T*_ that differ on some leaf edge. Let *f* and *f*′ be two of these assignment functions. Then there must be at least one leaf edge that has different images under *f* and *f*′. Suppose we call this leaf node *a* and the edge *l*_*a*_, then that *f* (*l*_*a*_) ≠ *f*′(*l*_*a*_). WLOG, assume *f*′(*l*_*a*_) *> f* (*l*_*a*_). Since *T* is unambiguous, *a* has at least one sibling. Call this node *b*. Also, there has to exist another leaf node *c*, not necessarily a sibling of *a*. Let the edges leading to *b* and *c* be *l*_*b*_ and *l*_*c*_ respectively. Let *d* and *d*′ be the distance from the parent node of *a* (and *b*) to *c* under *f* and *f*′ respectively. Then we have the following:

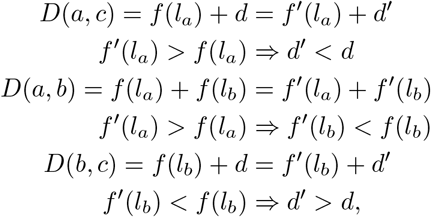

which is a contradiction. Therefore, all assignment functions agree at the leaf level. Now consider the subtree *T*′ derived from *T* by removing all leaf nodes. We can obtain 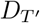 using 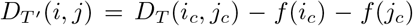 where *i*_*c*_ and *j*_*c*_ are any child node of *i* and *j* respectively and *f* is any assignment function. This is valid and will produce a unique 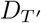 regardless of the choice of *i*_*c*_, *j*_*c*_ and assignment function because all assignment functions assign leaf edges to the same values and they all satisfy *D*_*T*_.

At this point, if *T*′ has only one level, then we can apply the special case and we are done. Otherwise, the same argument above can be applied to arrive at the conclusion that any assignment function for the edges of *T*′ agree at leaf level of *T*′. This procedure can be repeated by keep cutting away the leaves of the tree until we arrive at our special case, arriving at the conclusion that all assignment functions agree at every edge. In other words, the solution is unique.

The above result also implies that *A* is full rank.

Upon showing the existence of the solution, we now provide a simple algorithm to obtain this solution. To guide the understanding of the intuition behind the algorithm, we begin by making some observations with the aid of an illustration shown in Figure 1. Based on this illustration, we have two observations.

**Fig. 1:**
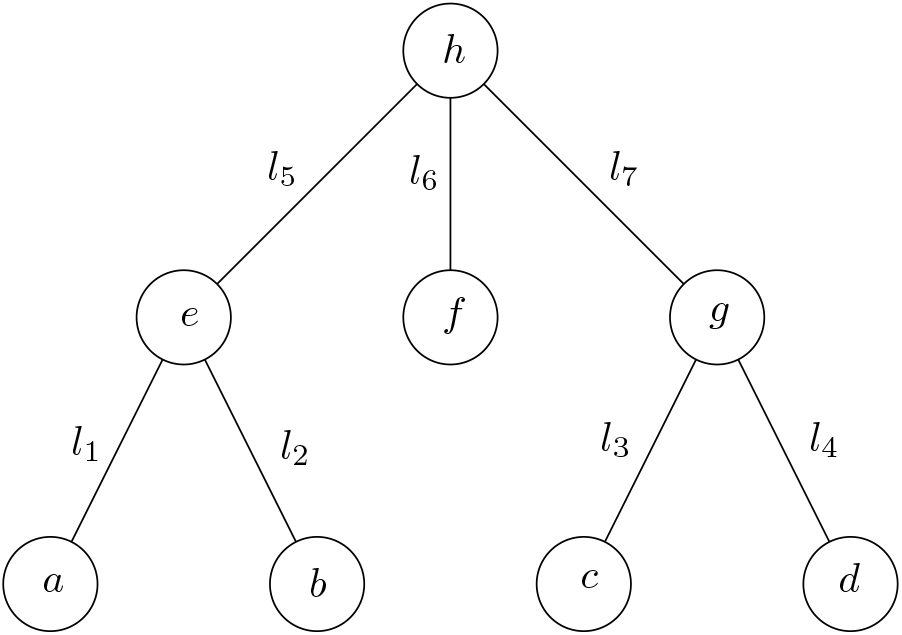
An example of an unambiguous tree for illustrative purpose. The label on each edge indicates the length of the edge.

#### Observation 1.

The lengths of all leaf edges can be recovered. For instance, *l*_1_ and *l*_2_ can be recovered as follows.

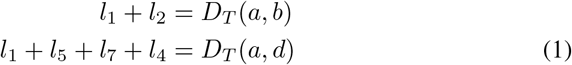

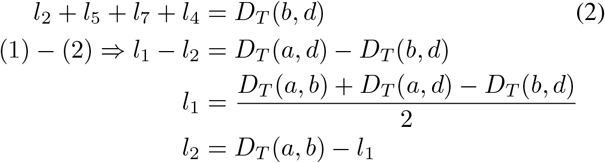

This procedure can be applied to every pair of siblings among the leaf nodes. Since *T* is unambiguous, every leaf node has a sibling and the additional leaf node (*d* in the case above) is always present.

#### Observation 2.

Given pairwise distances among nodes at level *k*, the distance matrix for nodes at level *k* − 1 can be computed.

Using Observation 1, we can compute all edge lengths at level *k*. Then, given any pair of nodes *i* and *j* at level *k* − 1, we can identify their respective children *i*_*c*_ and *j*_*c*_ at level *k*, of which pairwise distances are known. Then the distance between *i* and *j* can be computed by subtracting the the lengths of edges (*i, i*_*c*_) and (*j, j*_*c*_) from the distance between *i* and *j*. For instance, the distance between *e* and *g* in the illustration is *D*_*T*_ (*a, c*) − *l*_1_ − *l*_3_.

Combining these two observations, one can then successively compute the lengths of a level, followed by computing the pairwise distances of the level above, from the leaf level all the way up the tree until all the lengths of all edges are computed.

While this principle of solving branch lengths by grouping branches into triplets was previously conceived by Farris et al. [13], their objective was to use the triplets as guides to resolve tree topology instead of solving the branch lengths of a fixed tree topology. Furthermore, the description of this procedure was simply a narration of the principle, without rigorous and practical implementation.

In fact, despite the simplicity in narration, the actual implementation of the above-mentioned procedure can be inefficient in practice. Pairwise computation of distances at every level can be computationally intensive when the tree is large, and a large portion of the computed values are redundant. Here, we propose an efficient algorithm consisting of two steps as presented in Algorithm 1 and Algorithm 2 respectively, with edges cases and details omitted. The first step involves a topdown search to pre-select the triplets necessary for computing the branch lengths. The second step solves the branch lengths bottom-up using only the pre-selected triplets from the first part. In this way, we are able to achieve linear time complexity with respect to the number of edges, as demonstrated later in Figure 6.

#### Algorithm 1 search-needed-pairs

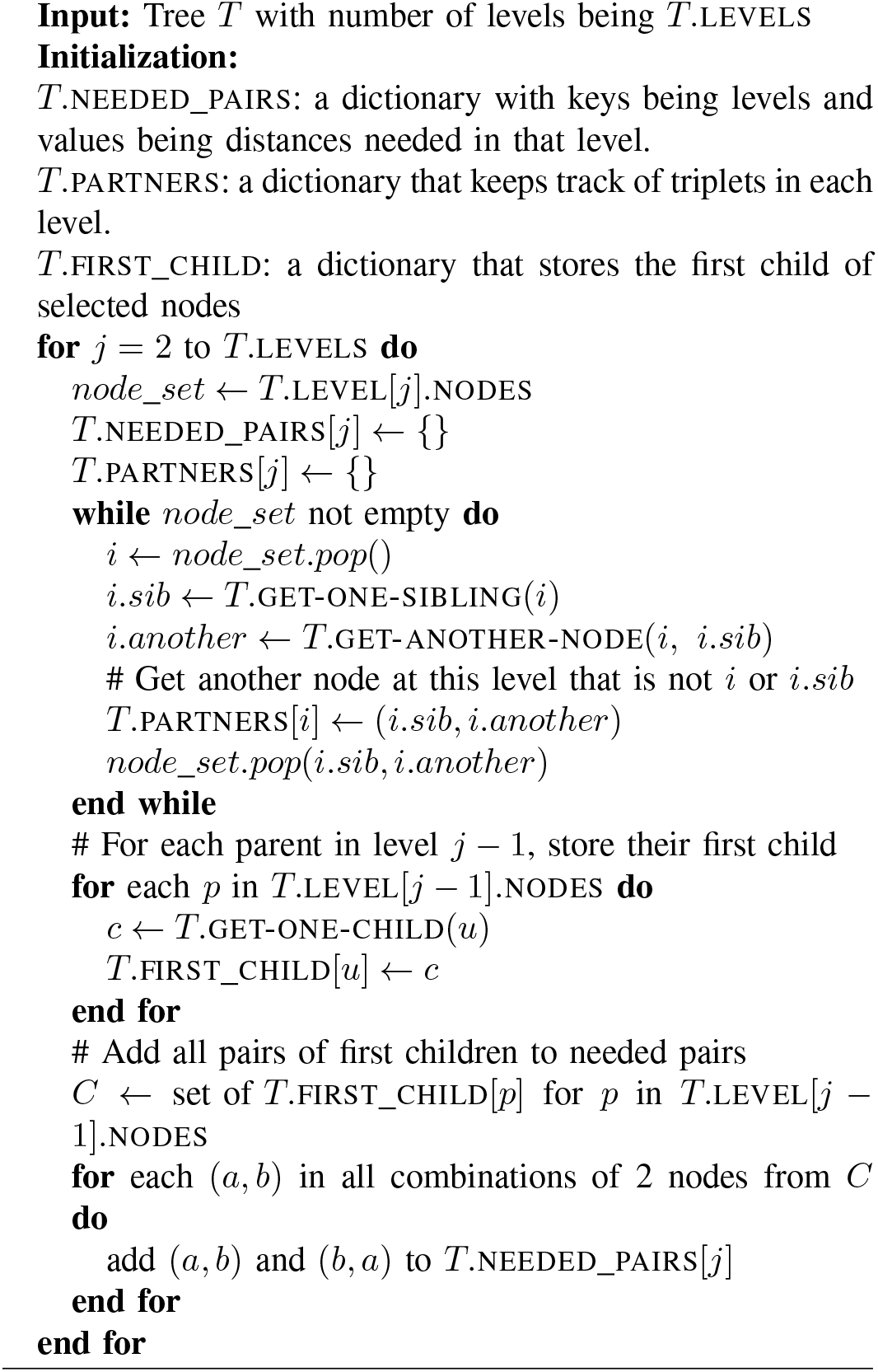

### C. A more realistic scenario

Algorithms 1 and 2 recover the edge lengths exactly when *T* is unambiguous and *D*_*T*_ compatible with *T*. In the real world, however, these two assumptions are often violated. For example, a binary tree is a commonly seen data structure in biological data, which is not an unambiguous tree. In addition, measurement errors due to limitations in equipment or measurement metrics are almost unavoidable, resulting in incompatible distance matrices. In the case of an ambiguous tree, there is nothing much that can be done without additional information. In the case of presence of single children or the root node having less than three child nodes, the exact lengths of the edges involved cannot be determined solely using the distance matrix. There are certain ways to mitigate this issue such as merging the edges connecting the single child, or manually assign a length that is biologically reasonable such as even splitting or the average length of edges in the same level. The exact handling of this issue may vary from situation to situation and there might not be a general way to measure which approach is superior to another. Therefore, this issue will not be discussed in depth in this paper and will instead be left to the discretion of researchers in specific situations.

#### Algorithm 2 solve-branch-lengths-bottom-up

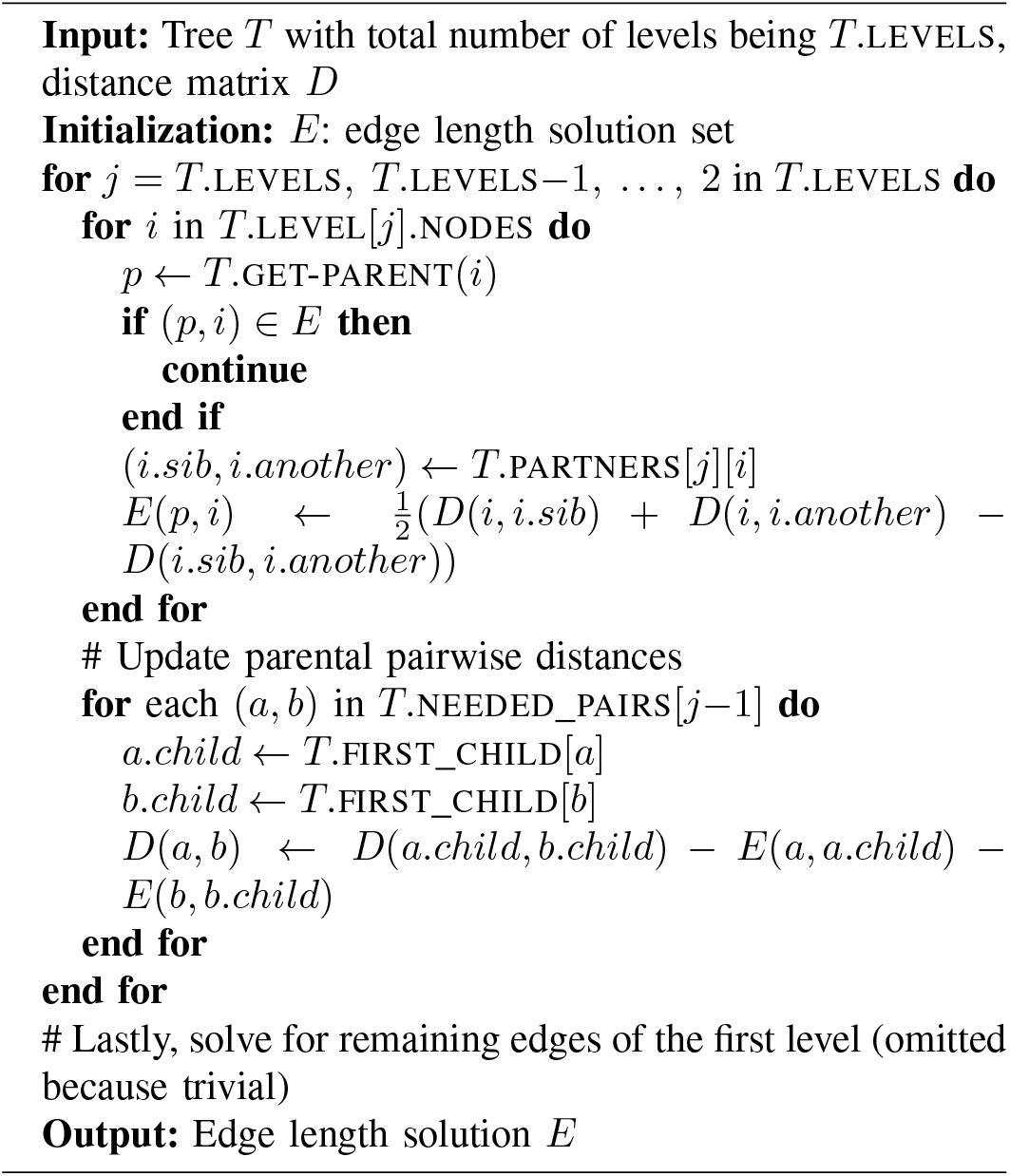

The second case with incompatible *D*_*T*_, on the other hand, is worth much more investigation as it is a problem that can be generalized. In the following discussion of this section, we will preserve the assumption that *T* is unambiguous while loosening the restriction that *D*_*T*_ is compatible with *T*. We modify the formulation of the problem in Section II-A as follows.

Suppose we are given an unambiguous tree *T* and an associated distance matrix *D*_*T*_, not necessarily compatible with *T*. Let 𝒟 represent the family of distance matrices associated with *T* that are compatible with *T*. Note that 𝒟 is not empty since one can always assign some lengths to the edges of *T* and compute pairwise distances among leaf nodes under this assignment, which will create a compatible matrix. Then, for any 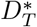 in 𝒟, there is a *y*^∗^ vector derived from 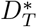 such that the linear system

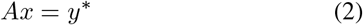

has a unique solution.

Let *y* be the vector derived from *D*_*T*_ and define *ϵ* = *y* − *y*^∗^. We can view *D*_*T*_ as a corrupted version of some 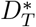 in 𝒟. In other words, *y* is some *y*^∗^ plus some error term *ϵ*. It is reasonable to assume that under normal circumstances where the measurement of *D*_*T*_ is reasonable and the incompatibility of *D*_*T*_ a result of minor errors, the difference between *D*_*T*_ and 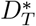 is small. As such, we pick 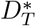 in D such that ||*ϵ*|| is the minimum and our objective now is to solve linear system while minimizing ||*y*^∗^ − *y*||. This equates to finding

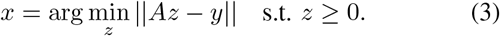

This can then be solved as a non-negative least square (NNLS) problem using existing algorithms and packages.

Alternatively, Algorithm 1 and 2 can be applied to give a solution, though it will not be consistent and will depend on the choice of *i*.*sib* and *i*.*another* in every iteration. To see this, consider the tree and distance matrix in Figure 2. If we compute *l*_*a*_ using nodes *a, b* and *c* we have

**Fig. 2:**
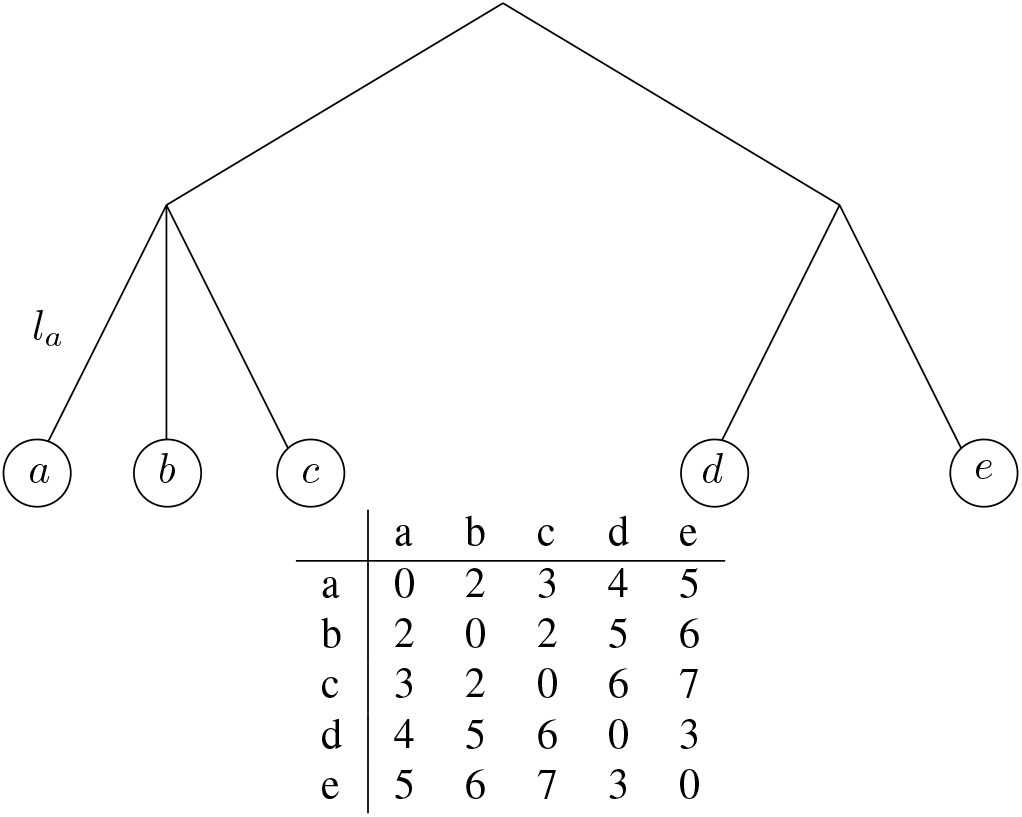
An example of a tree with an incompatible distance matrix.

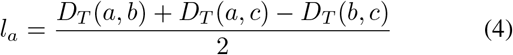

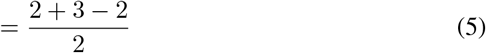

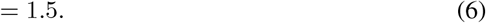

However, if we choose to use *a, c* and *e* instead, we have the following:

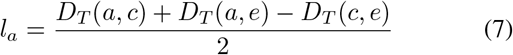

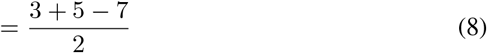

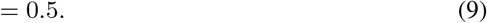

In general, if we let 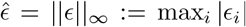, then at any leaf edge *l*_*a*_ with choices of sibling node *b, c* and choices of another node *d* and *e*, the “deviation” from *l*_*a*_ from the “true” length 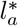 obtained by solving linear system (2) is given by

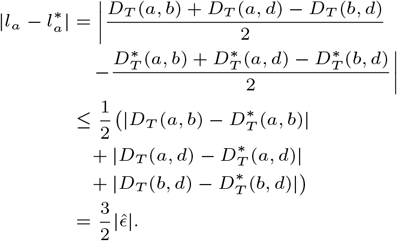

This bound is likely to increase up the tree as the “deviations” in edge lengths of one level will get carried over to the computation of pairwise distances among nodes of the parental level. Repeating the algorithm using different choices of *i*.*sib* and *i*.*another* and taking the average might help mitigate this issue. In the next section, we evaluate the performance of these methods in different scenarios using simulated data.

## III. Evaluation of methods using simulated data

In the previous section, we gave a brief outline of two potential methods that can be employed to solve linear system (2). In this section, we evaluated their respective performance using simulated data. To begin, we reiterate the methods we are comparing by giving each method a formal name and description.

### A. Method description

#### Definition III.1

(Naive NNLS). Given *A* and its associated *y*, the naive NNLS method solves the problem

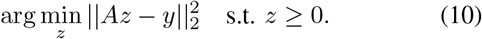

The prefix “naive” is to distinguish the above method from the “regularized NNLS” method described in the Supplementary Section 2, which can be considered a variation of the naive NNLS method. In our experiments with simulated data, the regularized NNLS method had a far worse performance than the other methods and hence will not be discussed in detail. The actual implementation of the naive NNLS solver is done using the `lsq_linear` function in the `scipy` package which employs the trust-region-reflect method of [17]. Since the matrix *A* has significantly more rows than columns, to enhance the efficiency of the computation, instead of using the entirety of *A*, we randomly select a portion of rows of *A* and use this smaller subset of *A* for the computation. The number of rows selected is tuned by a parameter *k* that represents a multiple of the rank of *A*. Since *A* is a full rank matrix, the rank of *A* is simply the number of columns of *A*. Since this implementation is randomized in nature, this procedure will be repeated a user-defined number of times and the average result of all the repeats will be taken as the final solution.

Alternatively, we define a second method as follows.

#### Definition III.2

(The Bottom-up method). This refers to solving for branch lengths bottom-up by selecting triplets of nodes as demonstrated in Algorithms 1 and 2. In the event that *D* is not consistent with the tree *T*, node-selection will be performed in random and the procedure repeated a user-defined number of times to produce an average solution.

In the next few sections, we evaluated the performance of these two methods from different aspects using simulated data. A glossary of the notations used in these experiments can be found in Table I.

**TABLE I:** Glossary of terminology used in Sections III-B through III-D.

| Notation | Name | Description |
| --- | --- | --- |
| $D_T$ | Distance matrix associated with $T$. | Matrix representing the pairwise distances among leaf nodes of $T$. |
| $A$ | $A$ | Coefficient matrix of the linear system $Ax = y$. |
| $y$ | $y$ | Measurement vector in linear system $Ax = y$, derived from $D_T$. |
| $k$ | Selection factor | $k \times \text{rank}(A)$ of rows will be used in the NNLS computation. |
| $p$ | Perturbation magnitude | The proportion of the distance that will be added to or subtracted from the original distance. |
| $s$ | Perturbation likelihood | The probability that perturbation will happen at any particular distance. |

### B. Experiment 1: recovering branch lengths using compatible distance matrices

We first assess the basic behaviors of these methods in the most ideal situation where *D*_*T*_ ‘s are compatible with the respective *T*. To this end we randomly generated *r*-ary trees with 1,000 nodes for *r* = 3, 5, 7, repeated ten times each. We assigned a random length between 0 and 1 to each of the branches and computed the associated *D*_*T*_ for each of the trees using the assigned branch lengths and used the two methods to recover the original assigned lengths using the computed *D*_*T*_. For the NNLS method, we set the selection factor *k* to be 5 and repeat number to be 100. For the bottom-up method, no repeat was needed since *D*_*T*_ ‘s are compatible.

The recovered branch lengths using each of these methods were compared against the original lengths and the correlation coefficient and *L*_1_ error were computed to quantitatively evaluate the performance of each method. Scatter plots of a selected portion of the results are also shown in Figure 3 to visualize the difference in performance among the three methods.

**Fig. 3:**
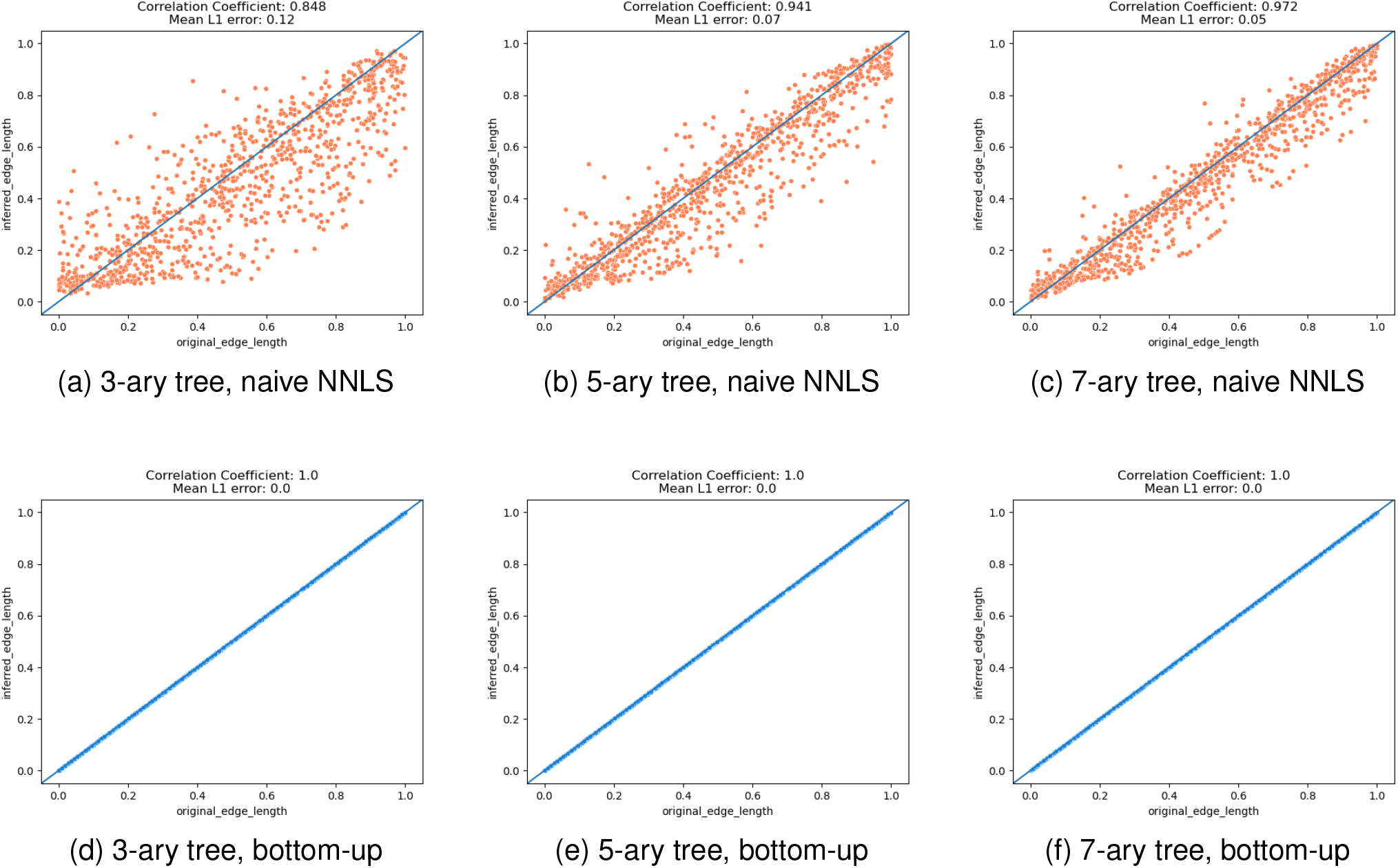
Scatter plots comparing the respective performance in recovering the original branches using different methods. Top: naive NNLS method. Bottom: bottom-up method. From left to right: 3-ary tree, 5-ary tree, 7-ary tree. X-axis: original edge lengths. Y-axis: recovered edge lengths.

As expected, in the case of compatible distance matrices, the bottom-up method perfectly recovered the original lengths while the naive NNLS method obtained solutions somewhat close to the original branch lengths, producing data points that formed a band approximately centered at the diagonal. The errors were most likely due to the fact that rows were selected at random during each iteration, which may not form a full-rank matrix, thus affecting the NNLS optimization procedure. To confirm this speculation and explore the effect of parameters, we repeated the same experiment using only the naive NNLS method, varying the selection factor *k* at different intervals from 5 to 200 while keeping the other parameters unchanged. The results are shown in Figure 4.

**Fig. 4:**
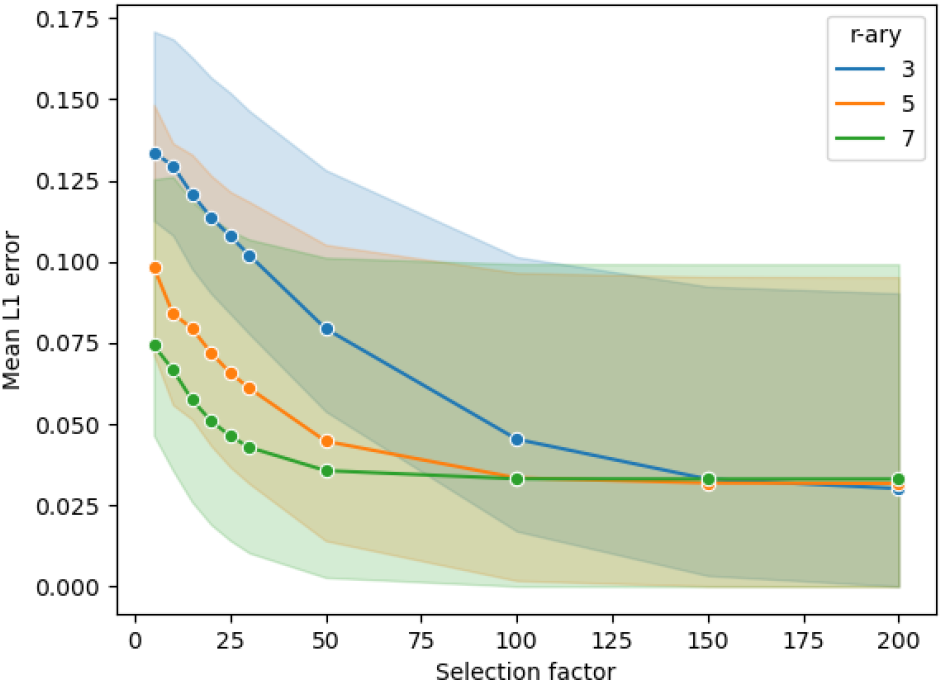
Mean *L*_1_ error between true branch lengths and those recovered by the naive NNLS method computed using different selection factors *k*, for 3-ary, 5-ary and 7-ary simulated trees each with 1,000 nodes.

As expected, *L*_1_ error decreased as more rows were used in the computation, though the time for computation also increased. Another interesting observation to note is that given the same number of nodes on *r*-ary trees, a higher value of *r* results in lower average *L*_1_ errors. This might suggest that the accuracy of the computation is sensitive to the height of the tree. As seen in Figure 4, a choice of anything above 50 might significantly reduce the error in our case. However, it is likely that this choice might not be generalizable to larger trees. The reason is as follows. For full 3-ary, 5-ary and 7-ary trees with 1,000 nodes each, the number of leaf nodes are 667, 800 and 857 respectively. The number of rows for matrix *A* for a given tree is the same as the length of vector *y*, which represents pairwise combinations of leaf nodes. Therefore, for full 3-ary, 5-ary and 7-ary trees with 1,000 nodes, the respective *A* matrix will have 222,111 rows, 319,600 rows and 366,796 rows. On the other hand, the rank of *A* is the number of edges, that is, 999, in all cases, since *A* is full rank. When the multiplication factor increases to 200, almost the entire matrix is used in the computation, hence achieving close-to-optimum results. However, when the tree size increases, the disparity between the number of rows and number of columns *A* increases exponentially. Hence, even 200 might not be the optimum choice. Ultimately, this problem boils down to a random covering problem. In subsequent experiments, we kept the original choice of 5 for the ease of computation, keeping in mind that increasing this multiplication factor will enhance the performance and tuning it as needed.

### C. Experiment 2: Recovering branch lengths using pairwise distances with errors

We now move on to the more complex scenario where the pairwise distances are not compatible with the tree. To simulate distance matrices that are incompatible with the tree, we introduced noise by perturbing the pairwise distances that are computed using the tree. To control the amount of perturbation, we introduced two parameters: perturbation magnitude 0 ≤*p* ≤1 and perturbation likelihood 0 ≤*s* ≤1. For each entry in *D*_*T*_, we generated a random number uniformly between 0 and 1. If the number was less than *s*, we randomly added or subtracted a proportion *p* of the original value, with equal probability. That is, *s* controls the proportion of *D*_*T*_ that was perturbed while *p* controls the magnitude of each perturbation. For example, if the original distance is *x*, then there is a 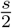 chance that the new branch length is *x* + *xp*, a 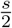 chance that it is *x* − *xp*, and a 1 − *s* chance that it remains *x*. We generated the same kind of trees as Experiment 1 for this experiment and set *p* to 0.01, 0.05, 0.1, 0.2, 0.4, 0.6 and 0.8 while setting *s* to 0.1, 0.2, 0.4, 0.5, 0.7 and 0.9. The settings for the two methods remained the same, except that we also repeated 100 times for the bottom-up method. The computed branch lengths were compared with the original unperturbed lengths and the *L*_1_ error was computed for each tree. Figure 5 shows the results.

**Fig. 5:**
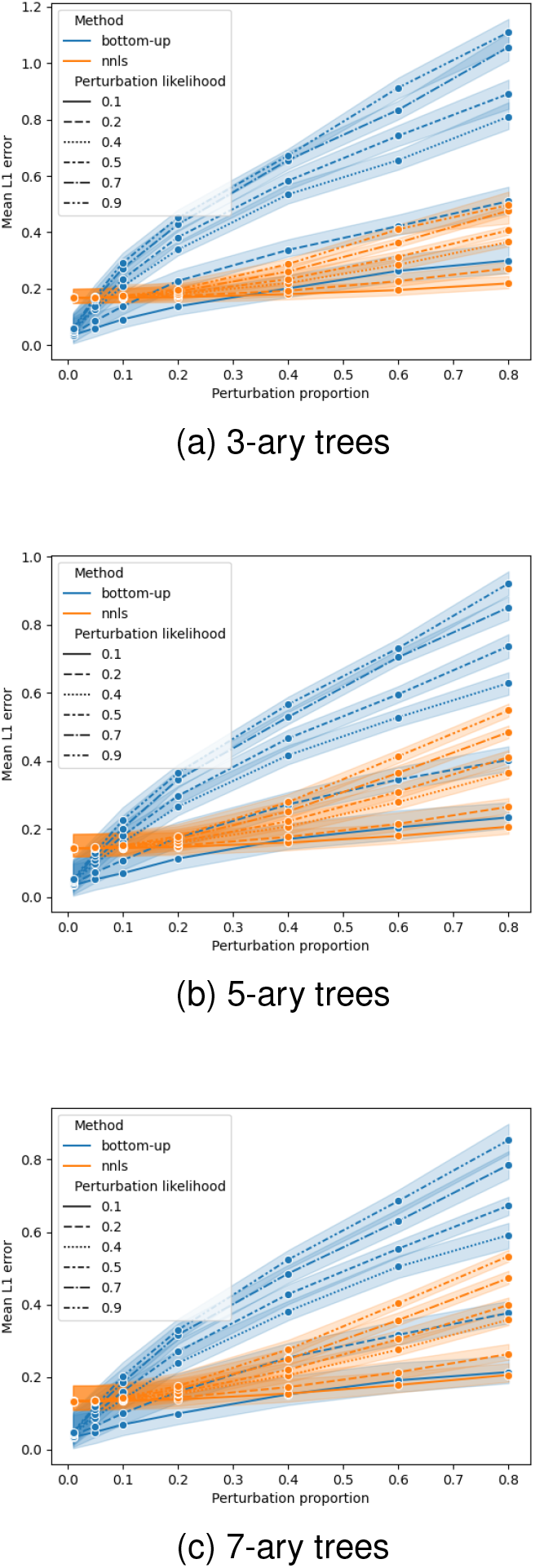
Robustness of methods against magnitude and frequency of perturbation, evaluated using mean *L*_1_ error. “Perturbation pro-portion” indicates the magnitude of perturbation introduced, while “perturbation likelihood” reflects the number of distances receiving the perturbation in the distance matrix. (a) 3-ary trees. (b) 5-ary trees. (c) 7-ary trees.

In general, the NNLS method has a comparatively better performance when the frequency of perturbation (perturbation likelihood) is high while the bottom-up method tends to perform better when perturbation is smaller. This suggests that in the event that the errors for distance measurement are known to be large, the NNLS method might be a better choice in terms of enhancing accuracy. An additional observation from the plots is that a similar trend to Figure 4 is observed here where 7-ary trees seemed to yield the lowest *L*_1_ errors on average, followed by 5-ary trees, with 3-ary trees yielding the highest average *L*_1_ errors, suggesting that accuracy may be inversely proportional to the number of levels.

### D. Experiment 3: Evaluating efficiency

In this section, we evaluate the performance of the two methods in terms of efficiency. To this end we generated 10-ary trees with increasing sizes, ten for each size, and computed the time needed for each of the methods. All distance matrices were perturbed with *p* = 0.4 and *s* = 0.5. To simplify the comparison, the process of repeating and taking average was omitted for both methods while all other parameters remained default. This was also to ensure the fairness of the comparison since there could be potential implementations that parallelize the repeating procedures to enhance the performance, which we will not discuss in detail in this paper. The result in Figure 6 shows the dominance of the bottom-up method in terms of speed. While running time for the NNLS method increased exponentially with increasing tree size, the running time for the bottom-up method remained almost constant. Theoretically, it can also be shown that Algorithms 1 and 2 runs in linear time. Additionally, the running time shown in Figure 6 did not include that of computing the *A* matrix. In real scenarios where the tree is large, the computation of matrix *A* alone may take up significant amount of time and memory, considering the dimension of *A*. In contrast, the bottom-up method uses only the pairwise distance matrix without computing and potentially storing intermediate data structures.

**Fig. 6:**
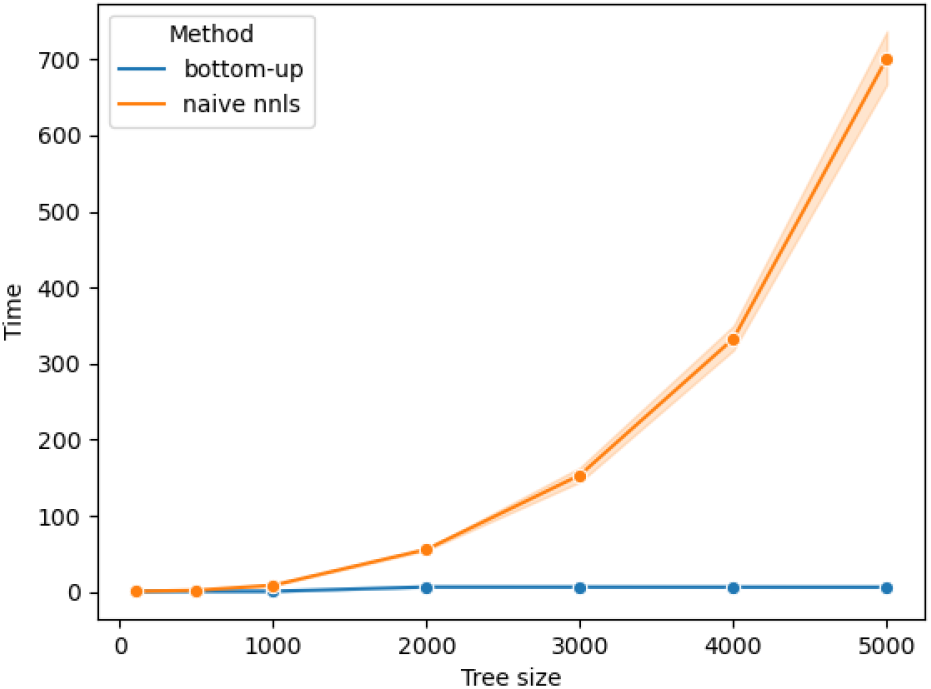
Running time for each method for solving trees with increasing size. X-axis: total number of nodes of the 10-ary tree. Y-axis: second.

### E. Summary

To summarize, the experiments using simulated data suggest that the NNLS method might be more robust against noises in the event where the distance matrix contains a high abundance of errors or inconsistencies, while the bottom-up method generally produces better results when noises are relatively small. In contrast, the bottom-up method works better when the distance matrix is close to a compatible matrix and is significantly superior in terms of efficiency. In the next section, we demonstrate a real-world application of branch lengths assignment of a tree with fixed topology while at the same time further investigate the performance of the two assignment methods in a real-life setting.

## IV. A biological application: FunUniFrac

One of the applications of imputing branch lengths of tree structures with fixed topology is that it extends the utility of tree metrics by allowing such computations to be performed on these tree structures. We demonstrate this by computing the “functional” UniFrac on the KEGG functional hierarchy tree [14]–[16], giving rise to a metric that measures the functional differences between metagenomic samples.

### A. Background

The UniFrac metric was originally designed as a metric to measure dissimilarity between metagenomic samples based on 16S rRNA data [8], providing insights to which organisms are differentially abundant in two metagenomic environments. This procedure involves three major steps. First, metagenomic samples are collected and represented as probability vectors indexed by discrete units such as the operational taxonomic units (OTUs) or amplicon sequence variants (ASVs). Second, obtain the phylogenetic tree with leaf nodes correspond to these units by either building it from scratch through tree-inferring algorithms or finding pre-built trees on databases such as GTDB [18]–[20]. Lastly, one computes the UniFrac distance between the OTU tables in the same way as the Earth Mover Distance over the tree [9].

This procedure can be adapted with the help of branch lengths assignment to answer a different yet equally meaningful question in metagenomic studies: the difference in functions that microbial communities are capable of performing in two given environments regardless of the actual organisms that carry out those functions. To do so, instead of clustering DNA into OTUs, we clustered them into functional units of orthologous genes through a process called functional profiling [21]. Next, instead of a phylogenetic tree, we obtained the KEGG Orthology (KO) hierarchy from the KEGG database [14]–[16]. The sources of this DAG consist of nodes labeled with “BRITE” ID, which stands for “Biomolecular Relationships in information Transmission and Exression”. We picked the first BRITE (ID ko00001) as the root and the spanning tree rooted at this node while removing others, such that we obtained a tree structure which we will call the “KEGG tree” henceforth. We computed pairwise the average amino acid identity (AAI) among the leaf nodes as a proxy for distance measurement by first converting KOs into sketches and followed by computing sketch similarity using `sourmash` [22], producing the distance matrix. Details for this procedure can be found in Section 3.1 in the Supplementary Materials. With this, we imputed edge lengths of the KEGG tree using both the NNLS method and the bottom-up method. A comparison between the lengths imputed using the two methods is shown in Figure 7. In general, the plot shows the proximity between the results obtained using the two methods, while suggesting that the NNLS method tend to produce smaller values in most cases.

**Fig. 7:**
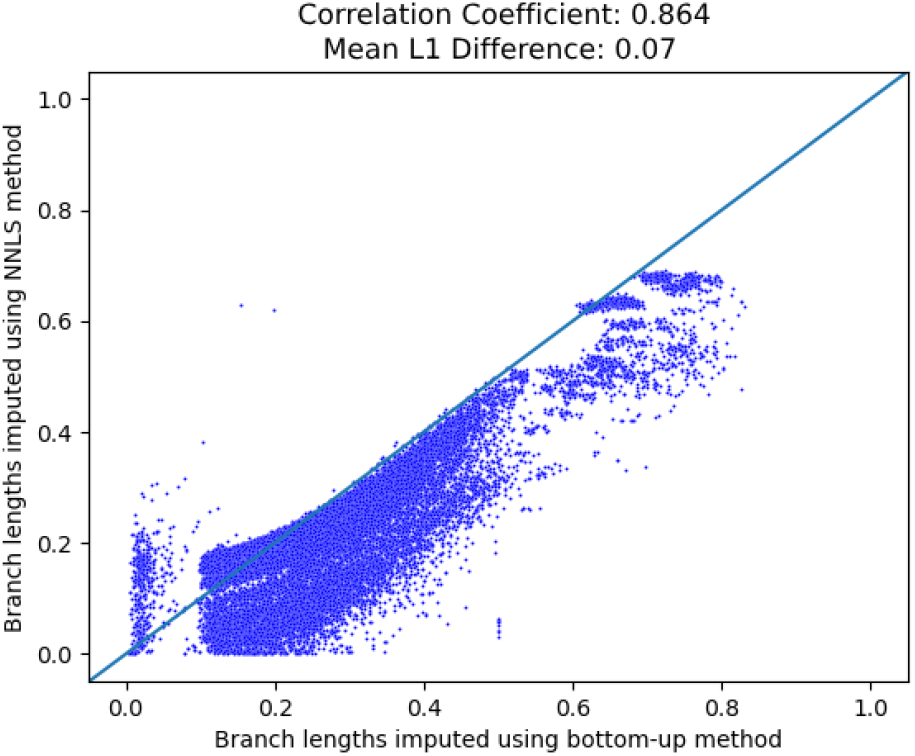
Comparison between branch lengths of KEGG tree imputed using the NNLS method and the bottom-up method. Each data point represents an edge, with the *x* coordinate being the length assigned using the bottom-up method and the *y* coordinate being the length assigned by the NNLS method.

As a further examination, we recomputed pairwise distances using both computed trees and compared the recomputed pairwise distances against the original pairwise AAI distances as well as against each other.

Figure 8 summarizes the comparisons of the recomputed pairwise distances. Overall, the pairwise distances computed using the two methods show similar trends, yielding a correlation coefficient of 0.979. In terms of the mean *L*_1_ error when compared to the original pairwise distances, the NNLS method seems to produce edge lengths more closely reflecting the original pairwise distances. In subsequent computations, we will use the “KEGG” tree with edge lengths imputed using the NNLS method, unless otherwise stated.

**Fig. 8:**
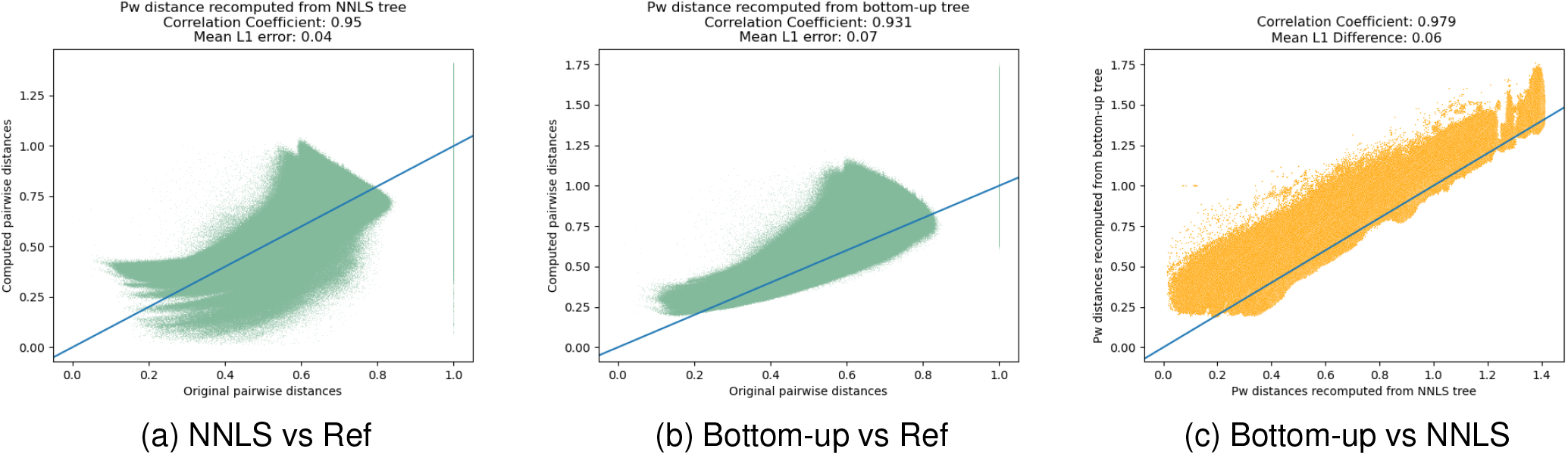
Analysis of recomputed pairwise distances from imputed trees. For 8(a) and 8(b): *x* axes represent the original pairwise *AAI* distances computed using `sourmash`; *y* axes: recomputed distances using the respective branch-lengths-imputed NNLS tree and bottom-up tree respectively by computing the tree distances. 8(c): a comparison between pairwise distances recomputed using the NNLS tree and the bottom-up tree. *x* axis: distance recomputed using the NNLS tree. *y* axis: distance recomputed using the bottom-up tree. The blue straight line across the window in all three plots serves as a guide to see how data points in each case deviate from the perfect case where the *x* coordinate equals exactly to the *y* coordinate for every point. The close the points are aligned with the blue straight line, the better the performance.

### B. Functional comparison among body sites

As a demonstration of this application, we downloaded metagenomic data from the HumanMetagenomeDB database [23] collected at four body sites: gut, oral, skin and vaginal, with 100, 72, 100 and 28 samples respectively. Each of these samples were converted into a functional profile in the form of a probability vector indexed by KOs, representing the abundances of the KOs in the sample. This is done using FracMinHash, [21] a sketch-based pipeline that uses `sourmash` methods to estimate the abundance of each KO present in each sample. The details for this process can be found in Supplement Section 3.2. Using the “KEGG” tree with imputed branch lengths, we computed the Earth Mover Distance between between any pair of samples over the tree, giving rise to a UniFrac-like metric measuring the dissimilarity between the samples in terms of protein functions. We term this metric the **FunUniFrac**, short for functional UniFrac. A Principle Coordinate Analysis (PCoA) was performed on the resulting pairwise FunUniFrac distance matrix and shown in Figure 9.

**Fig. 9:**
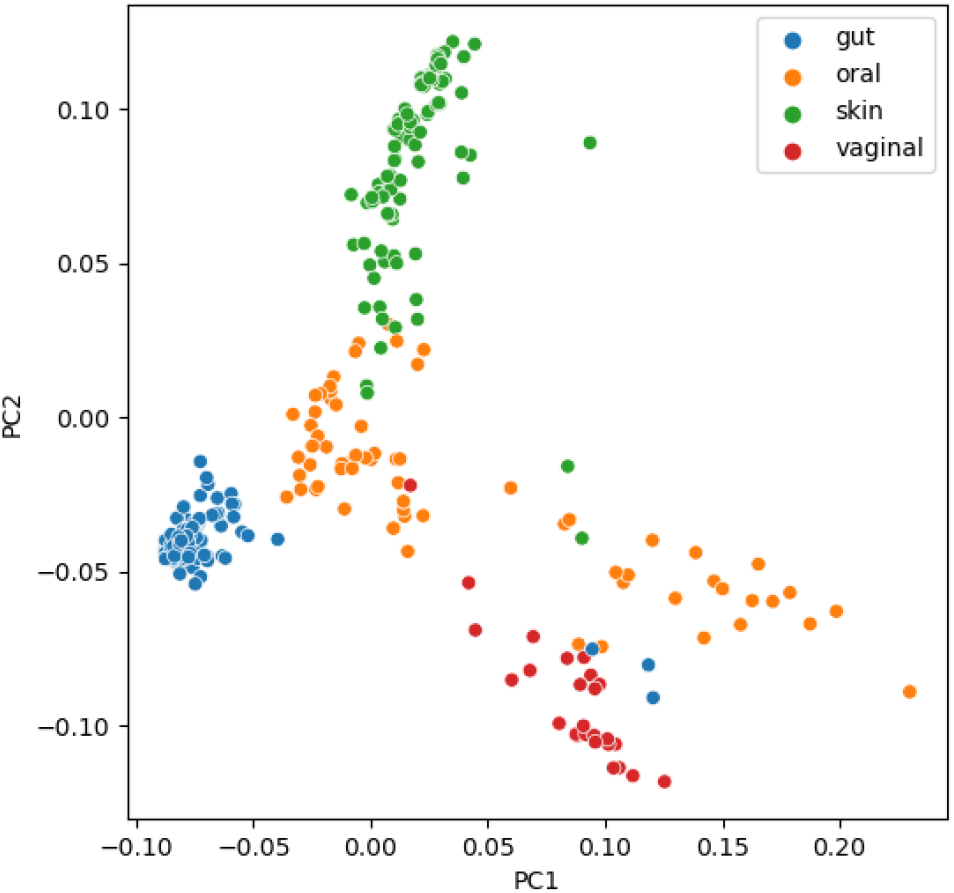
PCoA of 300 samples from four distinct body sites including gut, oral, skin and vaginal. Distances are based on FunUniFrac after functional profiles of the WGS metagenomes were obtained with the aid of FracMinHash.

**Fig. 10:**
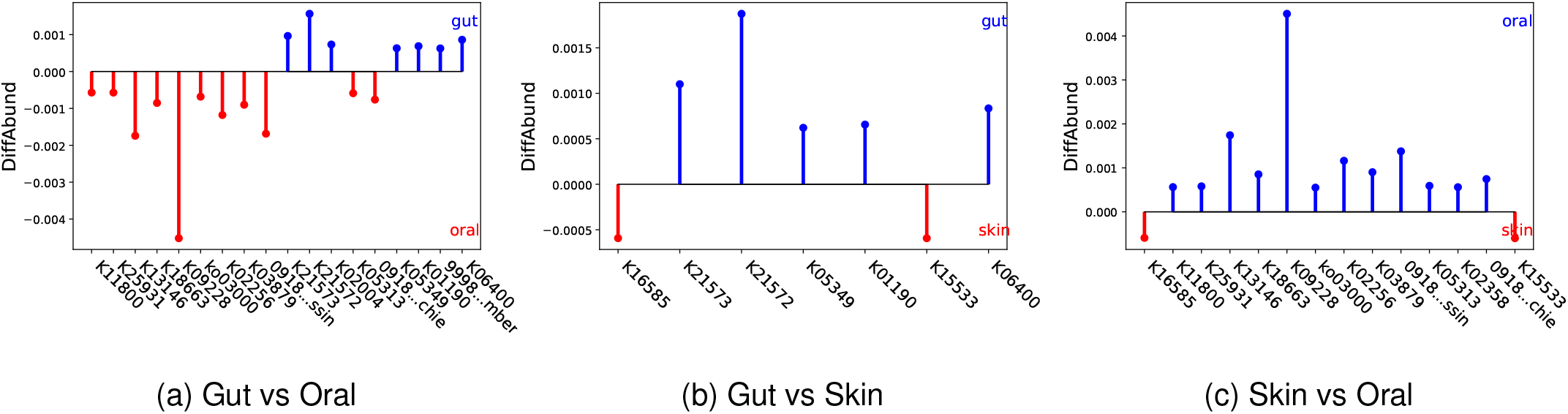
Pairwise “DiffAbund” vector among the average gut, oral, and skin samples. KEGG KO’s are given on the x-axis. Each vertical line indicates the amount of the relative abundance contributed by the corresponding KO to the FunUniFrac between the two environments being compared. The longer the line corresponding to a KO, the more the difference in relative abundance of the particular KO between the samples. A blue line indicates the KO is more abundant in the top-labeled sample and vice versa. Full function names of the KOs are listed in Supplementary Materials Table 3.

Judging from the PCoA plot, the samples are roughly clustered by body sites, recapitulating the trend of diversity of protein families across body sites shown in previous studies [24], [25]. This is also partially supported by the fact that metagenomic samples cluster by body sites in terms of taxa, as shown in Supplementary Figure 3. Interestingly, there are some distinct outliers in this plot. For instance, three gut samples are clustered distinctly away from the rest of the gut samples and are instead closer to oral and vaginal samples. Upon further investigation, these three gut samples were the only samples in our batch that were from one specific study of viromes [26] with viral DNA isolated and amplified. The fact that FunUniFrac revealed the two distinct clusters of gut samples suggests that there might be distinct functions performed by the viral proteins and that FunUniFrac metric is sensitive enough to capture these differences.

In Supplementary Section 5.2, we repeated the experiment using Bray-Curtis distance, a non-phylogenetic beta-diversity metric, and compared its performance against that of FunUniFrac. PERMANOVA test ruled in favor of FunUniFrac, showing the value of hierarchical information. Bray-Curtis also failed to capture the three outlier gut samples.

To further identify the exact functional proteins that significantly differ among these body sites, we obtained an average sample for each body site except for vaginal samples, which we excluded due to the fact that there were only a small number of samples, which may deem the average inaccurate by making the averaging process sensitive to outliers. To visualize the differences in the abundances of functional proteins, we computed what was called the “differential abundance (or DiffAbund) vector” [9] among the average samples for gut, oral and skin in a pairwise manner. The DiffAbund vector is a vector indexed by the functional units. Since the UniFrac is computed as an optimal “flow” between two samples and outputs a scalar value that quantifies the difference between two samples, the DiffAbund vector tracks the “flows” at any index in this process, thus quantifying the percentage contribution of each functional unit to the ultimate flow [9]. Through this process we were able to identify some KOs that were more prevalent in environment than others. For instance, K21572, named “starch-binding outer membrane protein” in KEGG database, is found in significant abundance in gut microbiome compared to skin samples. This agrees with past findings that gut microbiome are specialized for carbohydrates degradation [27]. On the other hand, K09228 is dominant in oral samples. According to the KEGG database, this KO bears the name “KRAB domain-containing zinc finger protein”, and was classified to be involved in the process of herpes simplex virus 1 infection [14]–[a virus type that causes lesions on mucus membranes, such as those in the oral cavity. Previous studies had also revealed that KAP1, a protein that interacts with the KRAB domain, was found to be significantly enriched in the presence of HSV-1 infection [28].

## V. Discussion

In this paper, we explored the problem of branch length reconstruction of trees with fixed topology given the pair-wise distance measurements of leaf nodes. Specifically, we proposed two methods, the NNLS method and the bottom-up method, that aim to solve this problem. We observed that these two methods each has its own strengths. The more the distance matrix is compatible with the tree topology, the better the performance of the bottom-up method becomes, while the NNLS method yields a more optimized solution when the distance matrix is highly incompatible with the tree. In terms of efficiency, however, the bottom-up method is highly superior over the NNLS method and can always be used as a guide when time constraint is present.

Obviously, there might be many other methods that can be utilized to solve this problem. It should be noted that the two methods discussed in this paper are presented not as the only two options but rather as representatives of two general classes of approaches for such problems. The NNLS method can be considered a global approach with the aim to achieve a global optimum guided by an objective function. The bottom-up method, on the other hand, does not take into account the global tree structure and focuses only on local solution at any point of computation. Each of these two methods can have many variations. For instance, different objective functions can be used for the NNLS method. Or perhaps, a regularization term may be incorporated into the objective function as shown in the Supplement. Though the regularized NNLS method is inferior to the naive NNLS method within the scope of this paper, the possibility that it could yield a better performance in some specific cases remains to be tested. For the bottom-up method, there are different options too. In our implementation, the edge lengths are solved bottom up starting from leaf level edges. Another alternative is to start from solving the lengths of longer paths, such as paths that start from a leaf and end at the root, using the same principle while treating a path as a long edge. By subtracting the lengths of shorter paths from the longer ones, the lengths of single edges can be solved from top down. Of course, in the event of an unambiguous tree coupled with a compatible distance matrix, these two methods will yield the exact same solution. However, in the event where these two conditions are not met, these two methods may favor different solutions. Another possibility is to combine the NNLS method and the local approach. The local methods, especially the “top-down” implementation, can be considered a form of divide and conquer approach in some sense. The experiments conducted in this paper showed that the NNLS method suffers from inefficiency but produces more globally optimal results when the distance matrix has a high degree of incompatibility. One can then divide the tree structure into smaller subtrees and solve each subtree using the NNLS method and “patch up” the solutions of these individual subtrees using the local approach to obtain a solution for the entire tree. This way, the efficiency can be significantly improved while maintaining some degree of sub-global maximality. Subsequently, the exact choice of approach is highly dependent on the nature of the data and research, and the methods presented in this paper are meant to serve as preliminary guidelines in making such choices.

Despite the straightforwardness of this problem, its application in computational biology should not be understated. In this paper, we demonstrated one of such applications by imputing the edge lengths of a protein function tree and thereby computing the “functional differential abundances” of metagenomic samples using this weighted functional tree as a basis. In some sense, this procedure creates a completely new metric (which we termed functional UniFrac) in metagenomic studies that allows researchers to analyze metagenomic samples from a new perspective and potentially draw new insights which have not been uncovered using traditional metrics. The KEGG protein function tree is only one of many such structures that bear the potential of benefiting from imputed branch lengths. Some other examples can include protein-protein interaction networks, metabolomic pathways, or any form of non-quantitative classification. It is important to note that both methods discussed in this paper are not limited to tree structures but can be applied to more general hierarchical structures such as forests, DAGs, or perhaps even general directed graphs. Further, a byproduct of branch lengths imputation is that it allows a biologically meaningful quantitative measurement to be computed among internal nodes, which are not naturally comparable numerically. For example, after imputing all the branch lengths for the KEGG tree, we can compute a distance between protein functions such as “methane metabolism” and “glycolysis” by subtracting the lengths of these nodes to their respective leaf-level descendants from the distance between those leaf nodes.

The main limitation or caveat in solving this problem lies in the choice of distance metric. In the case of leaf nodes with sequence representations, the alignment score could be a natural choice. In the case of the KEGG, we used AAI as a proxy for the distance between a pair of KOs. This choice is based on the nature of the representation of the KOs, i.e. orthologous proteins. The presence of multiple protein sequences in a KO makes alignment based measurement difficult, hence a *k*-mer based comparison was adopted instead. This choice is rather arbitrary and there can be other alternatives, such as decomposing KOs into motifs instead of k-mers, which might require a slight more effort. In general, the difficulty lies in finding a metric that best agrees with the topology of the tree or one that best fits the underlying biological question, and at the same time creates a distance matrix that is as compatible with the tree as possible, so as to give rise to the best biologically meaningful edge lengths. These conditions might be too stringent in most real-life cases and compromises might be inevitable, not to mention the fact that these qualities themselves might be hard to quantify due to a lack of ground truths. As such, the exact choice or design of distance metric may vary from case to case and remains an open area of research. Mathematically, it would also be helpful to come up with a way to measure the degree of compatibility of a distance matrix with a tree as well as methods or algorithms that corrects the distance matrix to maximize compatibility while minimizing the alteration.

## References

[1] W. M. Fitch and E. Margoliash, “Construction of phylogenetic trees: a method based on mutation distances as estimated from cytochrome c sequences is of general applicability.” Science, vol. 155, no. 3760, pp. 279–284, 1967.

[2] H. H. Otu and K. Sayood, “A new sequence distance measure for phylogenetic tree construction,” Bioinformatics, vol. 19, no. 16, p. 2122–2130, 2003.

[3] J. Rizzo and E. C. Rouchka, “Review of phylogenetic tree construction,” University of Louisville Bioinformatics Laboratory Technical Report Series, vol. 1, pp. 1–7, 2007.

[4] A. Rzhetsky and M. Nei, “A simple method for estimating and testing minimum-evolution trees,” Molecular Biology and Evolution, vol. 9, no. 5, p. 945, 1992.

[5] N. Saitou and M. Nei, “The neighbor-joining method: a new method for reconstructing phylogenetic trees.” Molecular Biology and Evolution, vol. 4, no. 4, p. 406–25, 1987.

[6] J. Felsenstein, “Evolutionary trees from dna sequences: A maximum likelihood approach,” Journal of Molecular Evolution, vol. 17, no. 6, p. 368–376, 1981.

[7] W. M. Fitch, “Toward defining the course of evolution: Minimum change for a specific tree topology,” Systematic Biology, vol. 20, no. 4, p. 406–416, 1971, maximum parsimony.

[8] C. Lozupone and R. Knight, “Unifrac: a new phylogenetic method for comparing microbial communities,” Applied and Environmental Microbiology, vol. 71, no. 12, p. 8228–8235, 2005.

[9] J. McClelland and D. Koslicki, “Emdunifrac: exact linear time computation of the unifrac metric and identification of differentially abundant organisms,” Journal of Mathematical Biology, vol. 77, no. 4, p. 935–949, 2018.

[10] G. D. Wu, J. Chen, C. Hoffmann, K. Bittinger, Y.-Y. Chen, S. A. Keilbaugh, M. Bewtra, D. Knights, W. A. Walters, R. Knight, R. Sinha, E. Gilroy, K. Gupta, R. Baldassano, L. Nessel, H. Li, F. D. Bushman, and J. D. Lewis, “Linking long-term dietary patterns with gut microbial enterotypes,” Science, vol. 334, no. 6052, p. 105–108, 2011.

[11] W. Wei and D. Koslicki, “Using the unifrac metric on whole genome shotgun data,” bioRxiv, p. 2022.01.17.476629, 2022.

[12] F. Pardi and O. Gascuel, “Combinatorics of distance-based tree inference,” Proceedings of the National Academy of Sciences, vol. 109, no. 41, p. 16443–16448, 2012.

[13] J. S. Farris, “Estimating phylogenetic trees from distance matrices,” The American Naturalist, vol. 106, no. 951, pp. 645–668, 1972.

[14] M. Kanehisa and S. Goto, “Kegg: Kyoto encyclopedia of genes and genomes,” Nucleic Acids Research, vol. 28, no. 1, p. 27–30, 2000.

[15] M. Kanehisa, “Toward understanding the origin and evolution of cellular organisms,” Protein Science, vol. 28, no. 11, p. 1947–1951, 2019.

[16] M. Kanehisa, M. Furumichi, Y. Sato, M. Kawashima, and M. Ishiguro-Watanabe, “Kegg for taxonomy-based analysis of pathways and genomes,” Nucleic Acids Research, vol. 51, no. D1, p. D587–D592, 2022.

[17] M. A. Branch, T. F. Coleman, and Y. Li, “A subspace, interior, and conjugate gradient method for large-scale bound-constrained minimization problems,” SIAM Journal on Scientific Computing, vol. 21, no. 1, pp. 1–23, 1999.

[18] D. H. Parks, M. Chuvochina, P.-A. Chaumeil, C. Rinke, A. J. Mussig, and P. Hugenholtz, “A complete domain-to-species taxonomy for bacteria and archaea,” Nature Biotechnology, vol. 38, no. 9, p. 1079–1086, 2020.

[19] “A standardized archaeal taxonomy for the genome taxonomy database,” vol. 6, p. 946–959, 2021.

[20] D. H. Parks, M. Chuvochina, C. Rinke, A. J. Mussig, P.-A. Chaumeil, and P. Hugenholtz, “Gtdb: an ongoing census of bacterial and archaeal diversity through a phylogenetically consistent, rank normalized and complete genome-based taxonomy,” Nucleic Acids Research, vol. 50, no. D1, p. D785–D794, 2021.

[21] M. R. Hera, S. Liu, W. Wei, J. S. Rodriguez, C. Ma, and D. Koslicki, “Fast, lightweight, and accurate metagenomic functional profiling using fracminhash sketches,” bioRxiv, p. 2023.11.06.565843, 2024.

[22] C. T. Brown and L. Irber, “sourmash: a library for minhash sketching of dna,” The Journal of Open Source Software, vol. 1, no. 5, p. 27, 2016.

[23] J. C. Kasmanas, A. Bartholomäus, F. B. Corrêa, T. Tal, N. Jehmlich, G. Herberth, M. von Bergen, P. F. Stadler, A. Carvalho, and U. Nunes da Rocha, “Humanmetagenomedb: a public repository of curated and standardized metadata for human metagenomes,” Nucleic Acids Research, vol. 49, no. D1, pp. gkaa1031.#x2013;, 2020.

[24] H. et.al, “Structure, function and diversity of the healthy human microbiome,” Nature, vol. 486, no. 7402, p. 207–214, 2012.

[25] E. A. Grice, H. H. Kong, S. Conlan, C. B. Deming, J. Davis, A. C. Young, N. C. S. Program, G. G. Bouffard, R. W. Blakesley, P. R. Murray, E. D. Green, M. L. Turner, and J. A. Segre, “Topographical and temporal diversity of the human skin microbiome,” Science, vol. 324, no. 5931, p. 1190–1192, 2009.

[26] S. Minot, S. Grunberg, G. D. Wu, J. D. Lewis, and F. D. Bushman, “Hypervariable loci in the human gut virome,” Proceedings of the National Academy of Sciences, vol. 109, no. 10, p. 3962–3966, 2012.

[27] J. F. Wardman, R. K. Bains, P. Rahfeld, and S. G. Withers, “Carbohydrate-active enzymes (cazymes) in the gut microbiome,” Nature Reviews Microbiology, vol. 20, no. 9, p. 542–556, 2022.

[28] L. D’Aiuto, D. C. Bloom, J. N. Naciri, A. Smith, T. G. Edwards, L. McClain, J. A. Callio, M. Jessup, J. Wood, K. Chowdari, M. Demers, E. E. Abrahamson, M. D. Ikonomovic, L. Viggiano, R. D. Zio, S. Watkins, P. R. Kinchington, and V. L. Nimgaonkar, “Modeling herpes simplex virus 1 infections in human central nervous system neuronal cells using two- and three-dimensional cultures derived from induced pluripotent stem cells,” Journal of Virology, vol. 93, no. 9, pp. e00 111–19, 2019.

