## Supplementary Materials for "On branch lengths assignment methods for trees with fixed topology and related biological applications"

### 1 Additional mathematics

#### 1.1 Examples of compatible and incompatible distance matrix

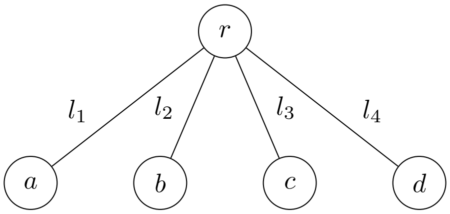


Figure 1: Tree $T$.

| | a | b | c | d |
|---|---|---|---|---|
| a | 0 | 3 | 4 | 5 |
| b | 3 | 0 | 5 | 6 |
| c | 4 | 5 | 0 | 7 |
| d | 5 | 6 | 7 | 0 |

Table 1: $D_T$: A pairwise distance matrix compatible with $T$.

| | a | b | c | d |
|---|---|---|---|---|
| a | 0 | 3 | 4 | 5 |
| b | 3 | 0 | 5 | 6 |
| c | 4 | 5 | 0 | 6 |
| d | 5 | 6 | 6 | 0 |

Table 2: $D'_T$: A pairwise distance matrix incompatible with $T$.

In this example, $D_T$ is compatible with $T$. Solving the equations gives a unique set of solution of $l_1 = 1, l_2 = 2, l_3 = 3, l_4 = 4$. $D'_T$ is an example of distance matrix that is not compatible with $T$. One way to see this is that on on hand we have $l_1 = \frac{1}{2}(D'_T(a,b) + D'_T(a,c) - D'T(b,c)) = \frac{1}{2}(3+4-5) = 1$. On the other hand, $l_1 = \frac{1}{2}(D'_T(a,c) + D'_T(a,d) - D'T(c,d)) = \frac{1}{2}(4+5-6) = 1.5$. These two solutions are inconsistent. In other words, there does not exist a mapping $f : E \to \mathbb{R}$ that agrees completely with $D'_T$.

#### 1.2 Additivity vs. compatibility

It is important to note that compatibility is different from the well-known concept of additivity and that a distance matrix can be additive and yet incompatible with an existing tree topology. To show this, we note that an $N \times N$ distance matrix is additive if and only if there is a metric tree with $N$ terminal nodes that generates the distance matrix [3]. Hence, the incompatible matrix $D'_T$ in the previous section is additive because there exists such a tree, shown in Figure 2.

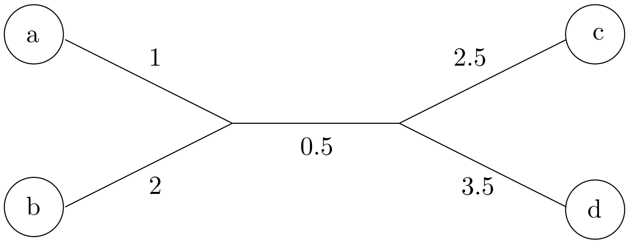


Figure 2: A tree that is compatible with distance matrix $D'_T$ in Section 1.1

### 2 Alternative method: regularized NNLS

An alternative method of solving the branch lengths problem besides the naive NNLS method and the bottom-up method is inspired by compressive sensing literature [2], which suggests the incorporation of a regularization term $\lambda||x||_1$ into the NNLS problem, giving rise to

$$\arg\min_x ||Ax - y||_2^2 + \lambda ||x||_1^2. \quad (1)$$

Combining the two norms, we have

$$\arg\min_x ||A'x - y'||_2^2 \; s.t. x \geq 0 \quad (2)$$

where $A' = [A; \sqrt{\lambda} ... \sqrt{\lambda}]$ and $y' = [y; 0]$. This constrained NNLS problem can be readily solved using existing algorithms.

We formally define the regularized NNLS method as below.

**Definition 2.1** (Regularized NNLS)**.** Given $A$ and its associated $y$, as well as a tunable regularization parameter $\lambda$, the regularized NNLS method solves the problem

$$\arg\min_x ||Ax - y||_2^2 + \lambda ||x||_1^2 \; s.t. x \geq 0. \quad (3)$$

When $\lambda$ is set to 0, this method is exactly the same as the naive NNLS method. This method was tested along with the other two using simulated data with compatible distance matrix in Section **??**, setting $\lambda$ to 1. The comparisons between the results are shown in Figure 3 below. As shown in the figures, the regularized NNLS method performed poorly compared to the other two. Decreasing $\lambda$ drove the performance of the regularized NNLS method to approach the performance of the naive NNLS method, suggesting that the naive NNLS method might be superior over the regularized NNLS method in general. As such, the regularized NNLS method was not included in the main manuscript.

### 3 Method details for FunUniFrac computation

#### 3.1 Computing leaf nodes distances

Upon downloading the tree structure of the "KEGG tree", we further downloaded the raw KO sequences from the KEGG website via FTP. These KOs are the leaf nodes of the "KEGG tree". As of November 2022, the downloaded data consists of 25,412 KOs and the entire tree has 25,865 nodes in total. After removing KOs with missing sequence data, the final KEGG tree consists of 25,524 nodes with 25,072 KOs as leaf nodes.

Each KO consists of one or more protein sequences. To compute the AAI distances among the KOs, we first sketched them using the "sourmash sketch" command of the `sourmash` package [1], setting the "kmer size" parameter to 5 and "scale factor" to 1,000. The KO sketches were then compared using the "sourmash compare" command, giving rise to pairwise distances.

#### 3.2 Converting metagenomic samples to functional profiles

To convert a metagenomic sample to functional profile, we first sketched the sample using "sourmash sketch translate" command of the `sourmash` package, setting the "kmer size" to 11 and "scale factor" to 1,000. The output of this step is an intermediate file containing signatures of proteins in this sample. We call this intermediate file the sample signature file. At the same time, we sketched the 25,072 KOs using the same parameters, giving rise to a database of signatures. Next, the sample signature file was searched against the signature database using the command "sourmash gather" from the `sourmash` package, giving us a functional profile of an estimate of the relative abundance of each KO contained in this sample.

### 4 Full KEGG function names

### 5 Additional comparison between FunUniFrac and other metrics

#### 5.1 PCoA based on WGSUniFrac distances

Figure 4 shows the PCoA plot of 300 metagenomic samples of four different body sites based on their WGSUniFrac distances as a contrast to Figure 9 in the main manuscript, which is based on FunUniFrac

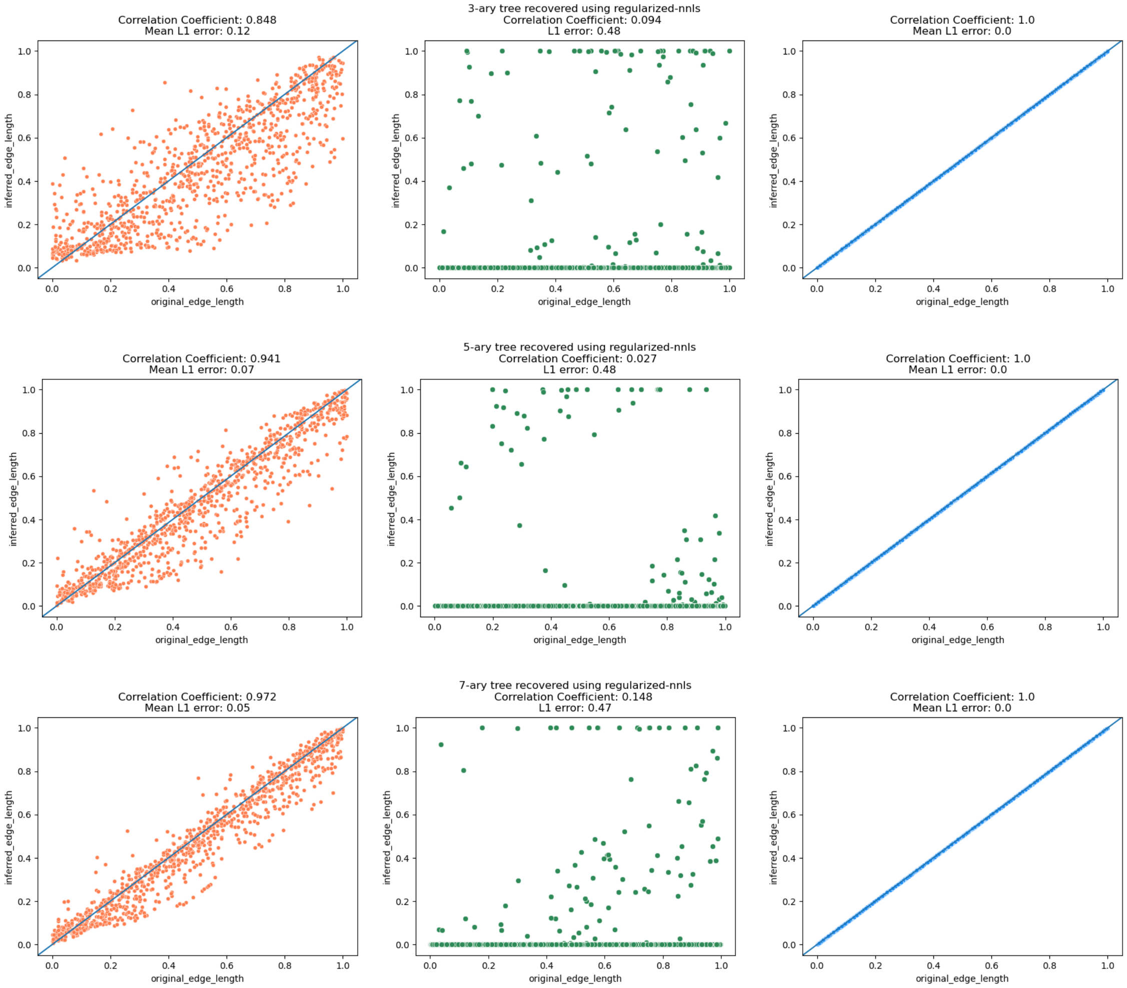


Figure 3: Scatter plots comparing the respective performance in recovering the original branches using the three methods. From left to right: naive NNLS (orange), regularized NNLS (green), bottom-up (blue). From top to bottom: 3-ary tree, 5-ary tree, 7-ary tree. X-axis: original edge lengths. Y-axis: recovered edge lengths.

distances.

The three labeled samples are the outliers in Figure 9 of the main manuscript. We notice that these three samples are not distinctively separated with the rest of the gut samples when clustered using WGSUniFrac, possibly due to limitations of the taxonomic profiling tool.

#### 5.2 FunUniFrac vs. Bray-Curtis

Figure 5 compares the PCoA plots of the same 300 samples from four body sites based on FunUniFrac distances and the non-phylogeny-aware Bray-Curtis distances.

While samples are generally clustered by body sites in both plots, clustering based on FunUniFrac is shown to have a higher quality as demonstrated using pseudo-F test, shown in Table 4. Additionally, the three outliers are not being distinctively captured using Bray-Curtis distance, suggesting the value hierarchical structure in functional comparison.

| Abbreviated name/KO | Full name |
|---|---|
| 0918...ssing | 09182 Protein families: genetic information processing |
| 0918...chies | 09180 Brite Hierarchies |
| 9998...mbers | 99980 Enzymes with EC numbers |
| ko03000 | Transcription factors |
| K01190 | Beta-galactosidase |
| K02004 | Putative ABC transport system permease protein |
| K02256 | Cytochrome c oxidase subunit 1 |
| K02358 | Elongation factor Tu |
| K03879 | NADH-ubiquinone oxidoreductase chain 2 |
| K05313 | Glutamate receptor, ionotropic, invertebrate |
| K05349 | Beta-glucosidase |
| K06400 | Site-specific DNA recombinase |
| K09228 | KRAB domain-containing zinc finger protein |
| K11800 | DDB1- and CUL4-associated factor 5 |
| K13146 | Integrator complex subunit 9 |
| K15533 | 1,3-beta-galactosyl-N-acetylhexosamine phosphorylase |
| K16585 | HAUS augmin-like complex subunit 2 |
| K18663 | Activating signal cointegrator complex subunit 3 |
| K21572 | Starch-binding outer membrane protein, SusD/RagB family |
| K21573 | TonB-dependent starch-binding outer membrane protein SusC |
| K25931 | NF-kappa-B-activating protein |

Table 3: Full names of all differentially abundant KOs and functions identified in Figure 10.

| **Metric** | **pseudo-F** | **p-value** |
|---|---|---|
| Bray-Curtis | 50.04 | 0.001 |
| FunUniFrac | 56.89 | 0.001 |

Table 4: PERMANOVA results comparing microbial community structure across body sites using Bray-Curtis and FunUniFrac distance metrics.

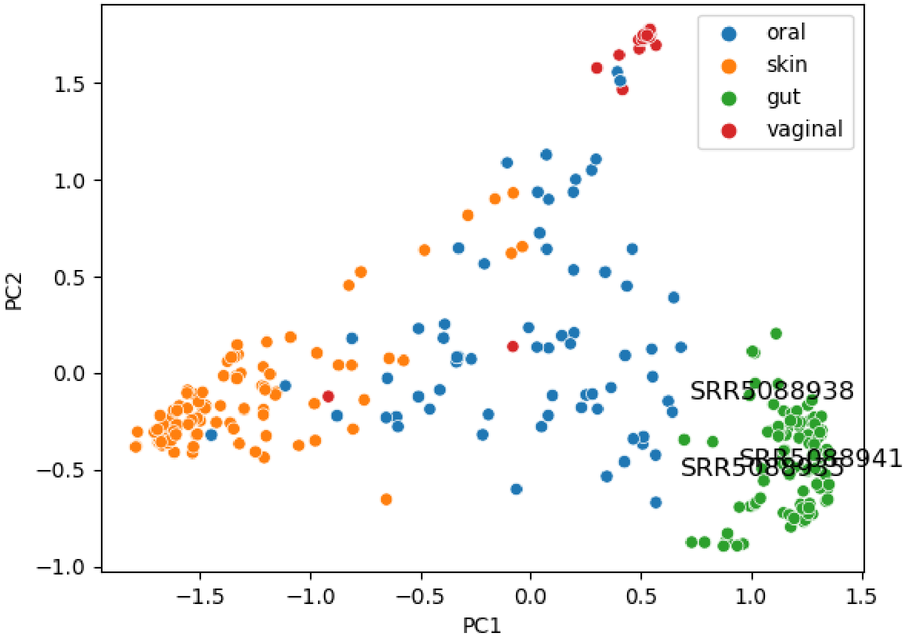


Figure 4: The PCoA plot of metagenomic samples of four different body sites based on their pairwise WGSUniFrac [4] distances. I.e. based on the proximity among their WGS profiles. Three of the gut samples are labeled as these three samples are shown as outliers in the PCoA plot in Figure 9 based on FunUniFrac distances. The three samples do not appear to be outliers in this plot. This could suggest that there might be some functional differences between these three samples and the rest of the gut samples, even though their taxonomic difference might not be significant enough to be captured in this PCoA plot. However, more studies are needed to arrive at a definite conclusion.

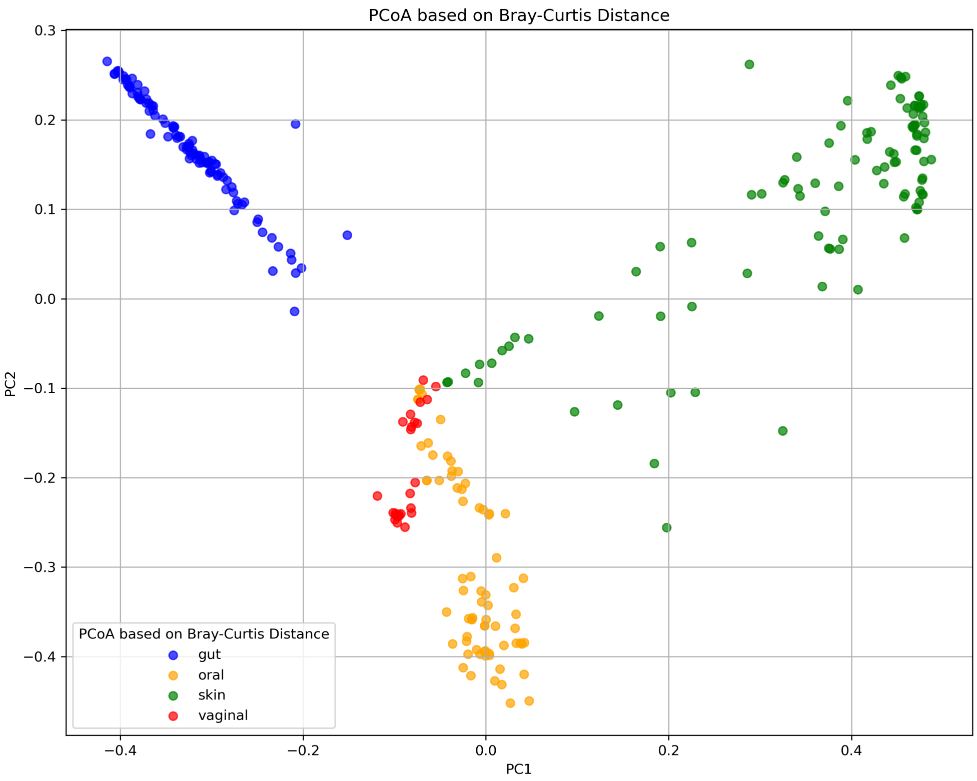


(a)

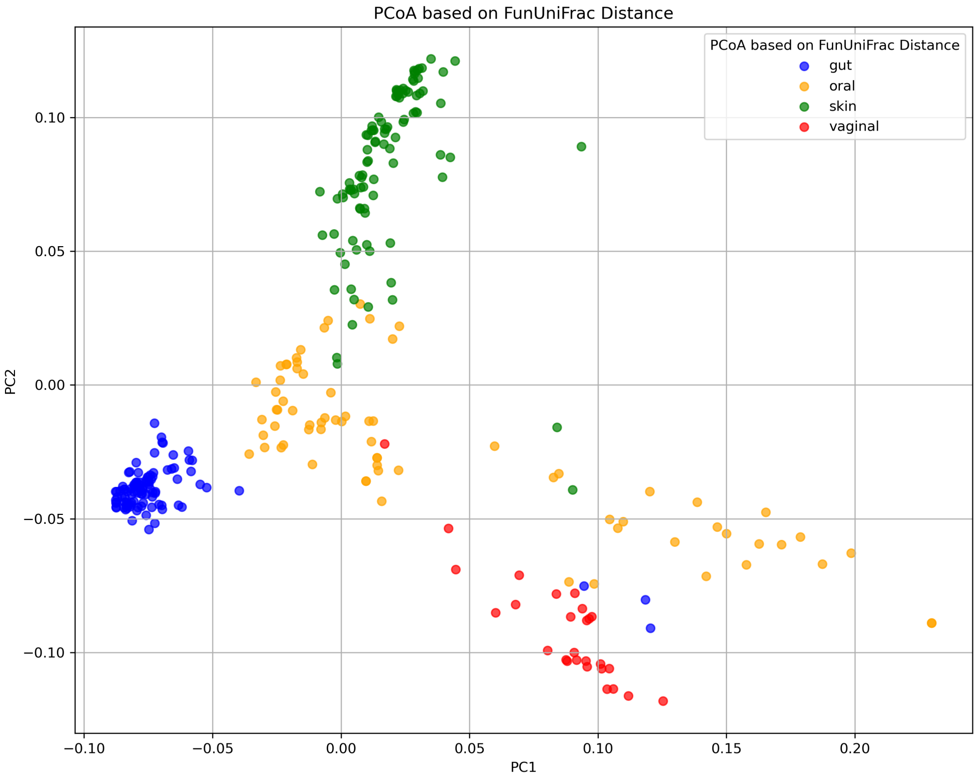


(b)

Figure 5: The PCoA plot of metagenomic samples of four different body sites based on pairwise Bray-Curtis distance (top) and pairwise FunUniFrac distance (bottom).

### References

[1] C. T. Brown and L. Irber. sourmash: a library for minhash sketching of dna. *The Journal of Open Source Software*, 1(5):27, 2016.

[2] S. Foucart and H. Rauhut. *A Mathematical Introduction to Compressive Sensing*, chapter 6. Birkhäuser, 2013.

[3] M. S. Waterman, T. F. Smith, M. Singh, and W. A. Beyer. Additive evolutionary trees. *Journal of theoretical Biology*, 64(2):199–213, 1977.

[4] W. Wei and D. Koslicki. Using the unifrac metric on whole genome shotgun data. *bioRxiv*, page 2022.01.17.476629, 2022.
